# Hierarchical binding of cognition to action in the human basal ganglia–thalamic circuit

**DOI:** 10.64898/2026.02.10.703559

**Authors:** Dennis London, Marisol Soula, Ling Pan, Michael Pourfar, Alon Mogilner, Roozbeh Kiani

## Abstract

Flexible behavior requires binding actions to their underlying cognitive variables, yet classical basal ganglia models emphasize a serial architecture where striatal action selection precedes a pallidothalamic motor gate. We tested this framework by recording single-neuron activity across the human pallidothalamic circuit while participants reported perceptual decisions with confidence-weighted reaching movements. Neurons in globus pallidus externus and internus, subthalamic nucleus, and motor thalamus exhibited sharp activity increases at movement onset that persisted through execution and feedback. Choice and confidence were continuously encoded before, during, and after movement. Population decoding revealed a transformation along the circuit, from dynamic upstream representations to stable, low-dimensional representations downstream, particularly in motor thalamus. After feedback, outcome signals emerged selectively in subthalamic nucleus and motor thalamus. In motor thalamus, confidence and outcome converged into a representation consistent with unsigned reward prediction error. These findings redefine the human pallidothalamic circuit as an interface that binds decision, action, and evaluation during behavior.

## Introduction

Cognitive flexibility allows us to adapt behavior to changing environments. Current theories often formulate decision-making as “thinking before acting”—a serial process in which decisions are made first, then actions follow (*1–3*). However, real-world behavior frequently requires cognition and movement to unfold and adapt in parallel. A midfielder advancing the ball down a soccer pitch exemplifies this dynamic, integrating team strategy, self-assessment of skill, and real-time environmental feedback while making precise movements. This ability to “think while acting” suggests that cognition and movement are not strictly sequential but dynamically interwoven in neural circuits (*4–6*). This kind of real-time control is simplest if evaluative variables are embedded within the same neural representations that specify the evolving action plan—an organization that would also support temporal credit assignment by keeping expectations and outcomes aligned with the actions that generated them (*7–9*).

Corticostriatal circuits are central to the computations that could support such integration. Neurons in the frontoparietal cortex encode decision formation (*10–13*), action plans (*14*), choice confidence (*15, 16*), action outcomes (*17–19*), and adjustment of behavioral strategies and decision policies based on past actions and outcomes (*20–22*). These signals converge in the striatum (Fig. 1A), which encodes sensory stimuli (*23*), perceptual decisions (*24, 25*), and expected reward (*26*), as well as the initiation, termination, kinematics, and sequence of movements (*27–32*). These observations place cognition and action signals in close anatomical proximity.

**Figure 1:**
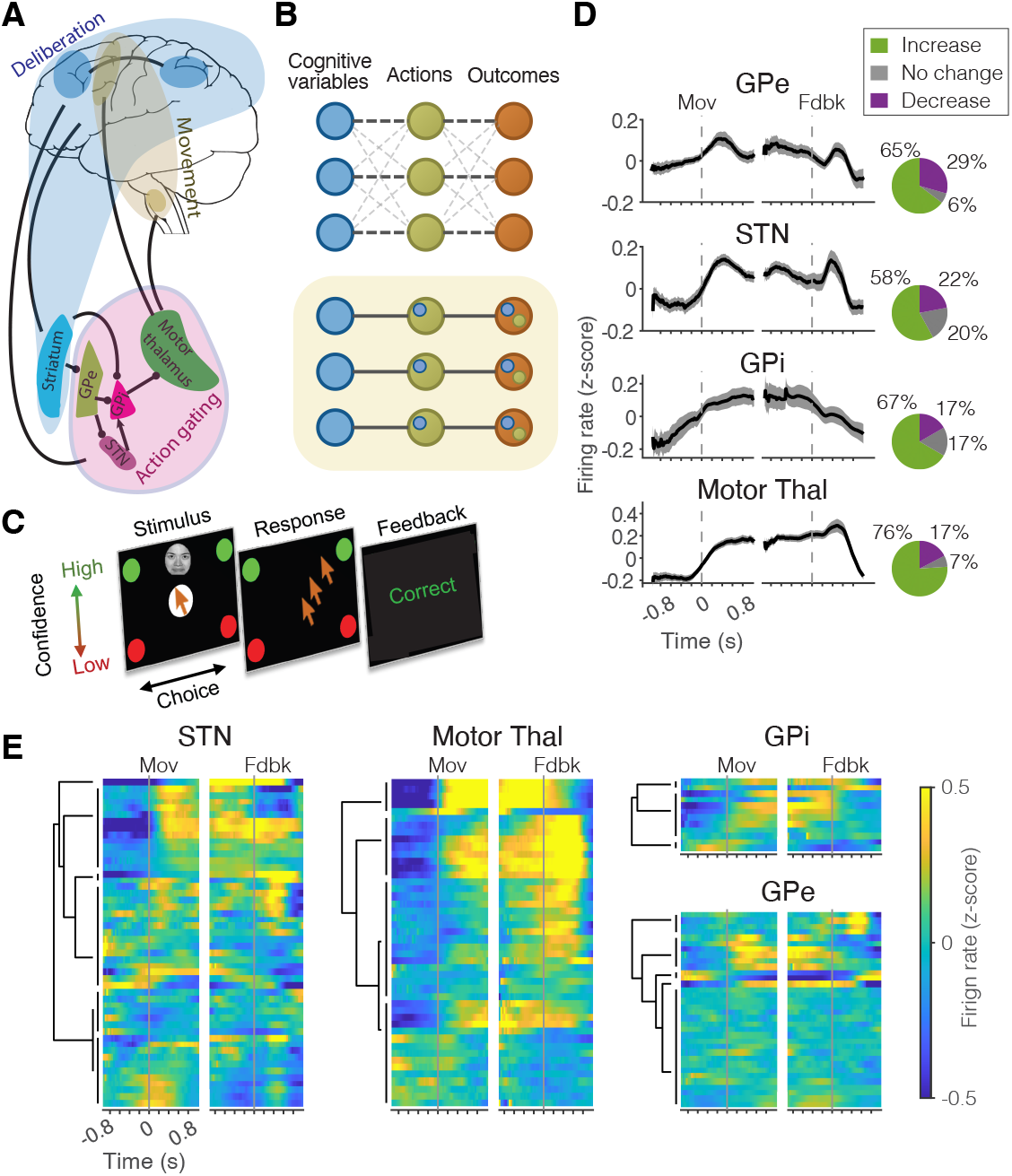
Persistent movement-linked firing rate increase across the human pallidothalamic circuit. (**A**) Current theories impose a separation between brain circuits involved in deliberation and movement, with the pallidothalamic regions implicated in gating and initiation of actions. (**B**) Flexible behavior requires cognition, action, and outcomes to be linked during ongoing behavior. We frame this as a cognition-action binding problem (top) and hypothesize a hierarchical solution where each action is co-encoded with its associated cognitive variables (bottom). (**C**) Perceptual decision-making task in which participants reported both choice and confidence using a single reaching movement. In different blocks, subjected discriminated facial expressions or motion direction of random-dot kinematograms. (**D**) Population-averaged peri-event time histograms aligned to movement onset and feedback in globus pallidus externus (GPe), subthalamic nucleus (STN), globus pallidus internus (GPi), and motor thalamus. Insets show the fraction of neurons with peri-movement firing rate increases. (**E**) Normalized firing rates of individual neurons aligned to movement onset and feedback. Neurons are sorted by similarity of their activity profiles, revealing functional clusters within each nucleus. Dendrograms were generated with hierarchical agglomerative clustering.

However, prevailing models of basal ganglia function posit a division of labor: corticostriatal circuits select among possible actions, while pallidothalamic circuits initiate the selected action by disinhibiting motor thalamus and downstream motor circuits (*33*). In these models, striatal medium spiny neurons convey action selection signals either directly to the output nodes of basal ganglia—globus pallidus internus (GPi) and substantia nigra pars reticulata (SNr)—or indirectly via the globus pallidus externus (GPe) and subthalamic nucleus (STN). The GPi and SNr, in turn, send inhibitory projections to the motor thalamus and brainstem motor centers, such that GPi/SNr suppression releases thalamocortical and brainstem circuits to execute the action. An implication of these models is that cognitive variables are progressively stripped away, yielding an action-gating “motor” code (*34–36*).

Several lines of evidence challenge key aspects of this feedforward framework. In primates, GPi activity shows weak or inconsistent correlations with motor thalamus (*37*), and GPi inactivation can paradoxically slow rather than facilitate movement (*30, 38*). In rodents, distinct functional cell populations in GPe and STN have been linked to both motor initiation and suppression (*39–43*). These results suggest a richer repertoire than a simple gate. In humans, how motor and cognitive signals are represented and transformed across pallidothalamic structures remains largely untested and unknown.

The human case is particularly important because our flexible behavior heavily depends on evaluating actions during execution and adjusting them based on context and outcomes. Perturbations of the STN and motor thalamus impair the adjustment of movements to feedback and parallel cognitive processes (*44–47*). Moreover, the human STN encodes ongoing movements and distinguishes planned from unplanned action adjustments (*48*), suggesting convergence of cognitive and motor information within this pathway. These findings motivate the hypothesis that pallidothalamic circuits do more than trigger movements; they may serve as an integral point between cognitive evaluation and motor execution during dynamic behavior. Here, we test this hypothesis by recording single-unit spiking activity across the human pallidothalamic pathway while subjects performed decision-making tasks that linked cognitive evaluation (confidence) with motor execution (reaching movements). This study presents the first direct recordings of spiking activity in the human pallidum during a cognitive-motor task, providing a critical test of action-gating theories of basal ganglia output. It also provides the first systematic recordings of spiking activity across human GPe, GPi, STN, and motor thalamus in the same cognitive-motor task, enabling us to ask when, where, and in what form cognitive variables become linked to action as behavior unfolds.

## Results

Twelve participants performed perceptual decision-making tasks where they categorized visual stimuli of varying degrees of ambiguity (difficulty) and made reaching movements toward one of four targets to report both their choice (perceived category) and their confidence level (high or low) (*49, 50*). Eight participants performed a stochastic face discrimination task (*51, 52*), judging facial expressions based on a dynamic stimulus. The other four subjects performed a motion discrimination task (*12, 20*) involving judgment of the net direction of motion of a field of moving dots (see Methods for task details). Despite differences in the visual stimulus, the tasks produced similar behavior and neural response dynamics, allowing us to pool the data (analyses of single task data matched the pooled data). To isolate motor-related activity, seven of these same subjects also performed a simple center-out reaching task, in which targets appeared in the same locations as the decision-making task targets. During task performance, we recorded single-neuron activity from GPe, GPi, STN, and motor thalamus. These recordings enable us to test whether the pallidothalamic circuit serves as a functional interface between decision-related cognitive processes and motor execution.

### Widespread pallidothalamic activation during movement initiation

We first test the predictions of classical prokinetic-antikinetic action initiation models of the pallidothalamic circuitry. These models posit that movement is facilitated by the direct pathway (prokinetic) and suppressed by the indirect pathway (antikinetic), leading to the predictions that neural response modulations in different pallidothalamic nuclei should (i) have opposite directions (e.g., increased activity in GPe and motor thalamus, decreased activity in STN and GPi), (ii) be relatively homogeneous within each nucleus, and (iii) occur primarily around the movement onset. An alternative hypothesis, largely untested in the human brain, is that diverse inputs from cortical and subcortical regions outside the basal ganglia critically contribute to activity in pallidothalamic nuclei, resulting in more heterogeneous, non-canonical response patterns.

Our data support this alternative. We observed that the onset of movement was accompanied by increased firing rates in all target nuclei—GPe, GPi, STN, and motor thalamus (Fig. 1, decision tasks and Fig. S1 instructed reaching task). The majority of neurons in each nucleus showed a sharp rise in firing rate aligned to movement onset (GPe: 65%, STN: 58%, GPi: 67%, motor thalamus: 76% of recorded neurons; Wilcoxon rank-sum test, p<0.05). In many neurons, this elevated activity persisted throughout the movement and extended into the feedback period (persistence defined as lasting for at least 75% of the movement epoch, see Methods; GPe: 64%, STN: 62%, GPi: 50%, motor thalamus: 80% of neurons with increased activity at movement onset; Wilcoxon rank sum test, p<0.05, Holm-Bonferroni corrected). In contrast, suppression of activity around movement onset was rare and typically transient, even within STN and GPi (STN: 22% of neurons showed a decrease, 10% persistent; GPi: 17% of neurons showed a decrease, 0% persistent). Finally, neural response dynamics in each nucleus were non-monolithic. Unsupervised clustering of activity profiles revealed multiple functional clusters with distinct activity dynamics (Fig. 1E), ruling out the notion of homogeneity within nuclei.

Importantly, these movement-related firing rate increases were observed across both the decision-making and instructed reaching tasks, indicating that they do not reflect a task-specific process (Fig. S1). Overall, these findings challenge the traditional interpretation of the prokinetic-antikinetic model and demonstrate that movement-related neural activity is excitatory and persistent with diverse dynamics throughout the pallidothalamic circuitry.

### Pallidothalamic encoding of cognitive variables before and during movement

We hypothesized that the diverse response patterns observed in the human pallidothalamic circuit arise, in part, from the integration of cognitive and motor information. To test this idea, we investigated whether non-motor variables could explain the observed firing patterns, focusing first on decision confidence. Confidence serves as a bridge between evaluation, action, and outcome, thereby facilitating learning and the adjustment of cognitive strategies (*3, 15, 49, 53*). A key design feature of our decision tasks was that subjects reported their choice and confidence using a single movement, ensuring confidence reports were not contaminated by post-choice processes. High confidence choices in our tasks were more likely to be correct and were associated with faster reaction times (Fig. 2A-B), consistent with past studies (*49, 50*). A significant fraction of neurons in the GPe (32%, p<0.001, binomial test), GPi (58%, p<0.001), STN (34%, p<0.001), and motor thalamus (20%, p<0.001) encoded confidence during movement (Figs. 2C-H and S2). In pallidum and STN, many confidence-encoding neurons began representing confidence during stimulus viewing (GPe, 58%; GPi, 43%; STN, 41%), more than 250 ms before movement initiation. In contrast, confidence encoding in motor thalamus emerged close to movement onset—no neuron encoded confidence more than 250 ms before movement onset (Fig. 2D; 0% selective earlier than 250 ms; 50% became selective between movement onset and 250 ms prior).

**Figure 2:**
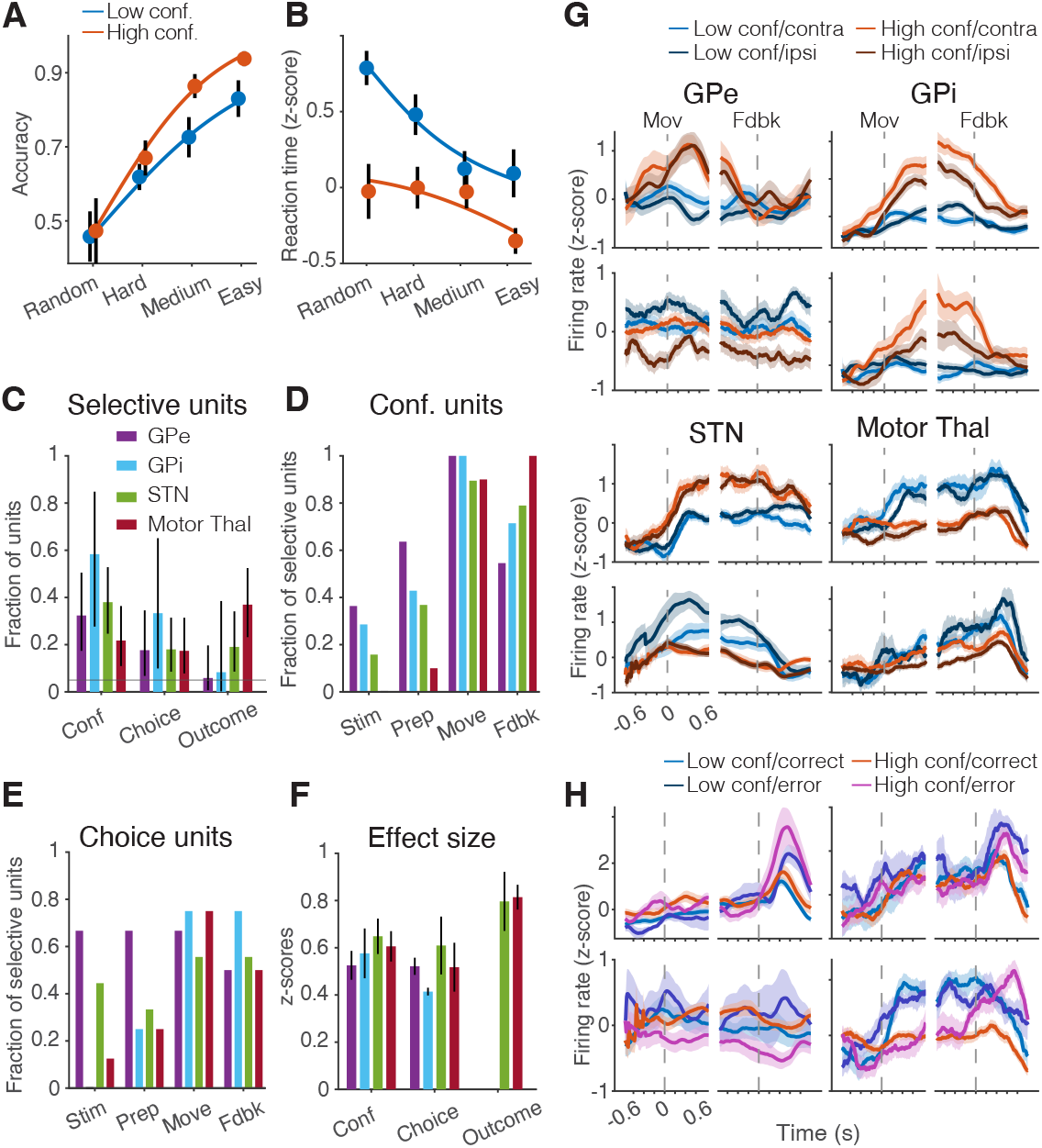
Single neurons encoded confidence, choice, and outcome with comparable strength, and increasing recruitment late in the trial. (**A**-**B**) Behavioral performance: high-confidence trials were associated with higher accuracy (A) and faster reaction times (B). (**C**) Fraction of neurons selective for confidence, choice, and outcome across task epochs in each region. Dashed line indicates chance-level selectivity; * indicates p<0.05. (**D-E**) Fraction of neurons selective for confidence (D) or choice (E) in each task epoch: stimulus viewing, movement preparation (250 ms before action onset), movement, and feedback. (**F**) Magnitude of firing rate modulation associated with confidence, choice, and outcome. (**G**) Example confidence- or choice-selective neurons from each pallidothalamic node. (**H**) Example outcome-selective neurons from STN (left) and motor thalamus (right).

Overall, these confidence signals emerge gradually during stimulus viewing, with the strongest representations during the movement period. Further, there is a systematic difference in the latency of these signals across areas, as we show here and elaborate in the following sections. These dynamics suggest a flow of confidence signals from the output nodes of basal ganglia to the motor thalamus.

Confidence-related firing rate modulations during the movement were robust (Fig. 2F), 33-106% greater than firing rate modulations for corresponding movement directions on the instructed reach task (Fig. S3, z-scored firing rate modulations: GPe, 0.52 vs 0.39, p=0.049; STN, 0.64 vs 0.31, p=0.008; GPi, 0.56 vs 0.39, p=0.03; motor thalamus, 0.60 vs 0.34, p=0.01). Further, the observed encoding of confidence was distinct from the encoding of movement and could not be attributed to a correlation between confidence and movement vigor (Fig. S4 and Supplementary Material) In pallidum and STN firing rates were highest for high confidence movements (GPe, 55%; GPi, 71%; STN: 69%), but, surprisingly, in motor thalamus *all* (100%) confidence-encoding neurons had the highest firing rates for low-confidence movements (example neurons in Fig. 2G), contrary to what would be expected if elevated movement vigor on high-confidence movements explained confidence-associated firing rate modulations.

Choice-related firing rate modulations on the decision tasks were present mainly during the movement itself and had a statistically similar magnitude as confidence-related activity (Fig. 2F; z-scored firing rate modulations: GPe, 0.52 vs 0.52, p=0.13; STN, 0.64 vs 0.62, p=0.25; GPi, 0.56 vs 0.42, p=0.5; motor thalamus, 0.60 vs 0.52, p=0.25). Further, the time course of the choice-related modulation was similar to modulation of firing rates with movement direction in the instructed reach task, indicating that, in contrast to confidence-related signals, choice-related signals encode the action rather than its underlying decision variables (Fig. S3).

Together, these findings challenge the traditional dichotomy between cognitive and motor processing in basal ganglia circuits. The former is typically studied while subjects are deliberating and before they make a response. In contrast, motor neuroscience focuses on the time period during movement and just before. These paradigms implicitly assume a model in which the neural computations or circuits that do the thinking are spatially and temporally distinct from those that do the acting. Instead, we find that neural populations in GPe, GPi, STN, and motor thalamus—structures implicated in gating selected actions—encode confidence dynamically and continuously before and during movement. Confidence-encoding persisted even after the completion of the movement and throughout the feedback period (GPe, 50% of neurons with confidence selectivity during movement; GPi, 71%; STN, 76%; motor thalamus, 100%). Furthermore, confidence is not the only encoded cognitive variable, as neurons in STN and motor thalamus also represented the outcome (correct vs. error) of the movement after feedback was delivered (fraction of all neurons: GPe, 6%, p=0.24; GPi, 0%, p=0.45; STN, 17%, p<0.001; motor thalamus, 35%, p<0.001; Fig. 2C), supporting a model in which the pallidothalamic circuit serves as a real-time interface between cognitive evaluation and motor output.

### Hierarchical binding of motor and cognitive signals in the pallidothalamic circuit

The joint encoding of movements and their underlying cognitive variables suggests a potential mechanism for solving the *binding problem*—the challenge of linking each of many simultaneously ongoing actions to its corresponding deliberative process (Fig. 1B). If the pallidothalamic circuit binds motor signals with associated cognitive variables at movement onset (e.g., choice confidence) it can broadcast these jointly encoded signals via the motor thalamus to widespread cortical and subcortical targets. This binding would enable “acting while deciding,” a hallmark of natural behavior (*4–6*).

For this function to be effective, motor and cognitive signals should meet three criteria: (i) they should be co-represented and linked, not independent, (ii) their representations should persist throughout the movement, and (iii) they should support stable readout throughout the movement, despite changes in movement kinematics. We hypothesized that if the pallidothamic circuit performs this motor-cognitive binding, these three characteristics should strengthen progressively along the circuit, from the upstream node (GPe) to the output node (motor thalamus).

We tested these predictions using a series of cross-temporal decoding analyses of population firing rates for different task variables. To define encoding axes in the neural population state space, we used linear regression to decompose firing rate modulations of each neuron at each time into components associated with choice, confidence, and outcome, as well as an intercept term. This intercept term captured changes of neural responses over time that were not explained by task variables and reflected the peri-movement rise of activity (see previous section). We projected neural population activity onto the encoding axis associated with each task variable defined by the vector of corresponding regression coefficients across neurons. The projections were well separated for the choices, confidence levels, and choice outcomes, allowing these variables to be decoded from population firing rates as quantified by a leave-two-out cross-validation procedure. On each iteration, we defined the decoding axes from n − 2 trials and predicted the corresponding task variable for the two held-out trials. We explored how these task encoding axes and their associated decoding accuracies evolved over time, both within and across trial epochs (Figs. 3 and S6).

**Figure 3:**
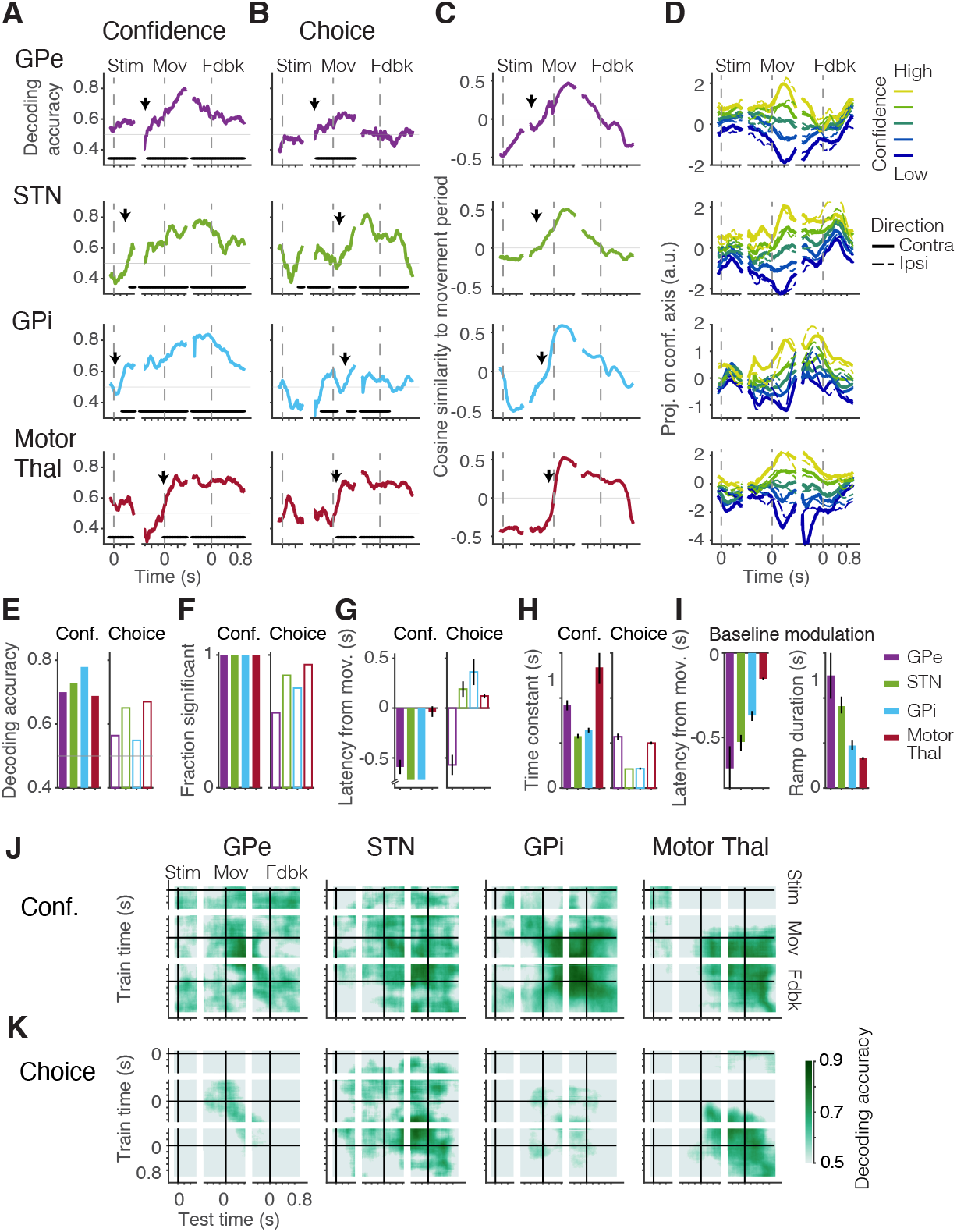
Hierarchical emergence and stabilization of cognitive and motor representations in the pallidothalamic circuit. (**A**-**B**) Decoding of confidence (A) and choice (B) over time using population encoding axes derived from population firing rates at each time point. Asterisks above abscissa indicate periods of above-chance decoding (cluster mass test, p<0.05). Arrows indicate the latency of above-chance decoding computed from piecewise exponential fits. (**C**) Cosine similarity of condition-independent (baseline) activity to movement-period activity (200 ms after movement onset). Movement and feedback period activity was anti-correlated with pre-movement baseline. Sigmoid fits were used to estimate the latency (arrows) and duration of this reversal of condition-independent activity. (**D**) Population activity stratified along the confidence-encoding axis (estimated from population firing rates 400 ms after movement onset), showing stable separation of confidence levels throughout movement. (**E**-**G**) Summary statistics: mean decoding accuracy during movement (E), fraction of the movement period with significant decoding of confidence and choice (F), and decoding latencies (G) (arrows in A-B). (**H**) Temporal stability of confidence and choice representations quantified with time constants of off-diagonal cross-temporal decoding in J-K (see Fig. S5). (**I**) Latency (arrows in C) and ramp duration of the reversal of condition-independent activity. (**J**-**K**) Cross-temporal decoding of confidence (J) and choice (K) revealed an increasingly stable code in downstream nodes of the pallidothalamic circuit. Decoders trained at one time point (y-axis) were used to decode activity at other time points (x-axis). Only statistically significant decoding is shown.

Confidence was reliably encoded by the neural populations in all nodes of the pallidothalamic circuit (mean decoding accuracy during movement: GPe, 70%; GPi, 79%; STN, 73%; motor thalamus, 68%). Confidence signals emerged >500 ms before movement onset in STN, GPe, and GPi (Fig. 3G). In stimulus-aligned analyses, confidence decoding became significant within 500 ms of stimulus onset in STN and GPi, with onset latencies comparable to or later than those reported in frontoparietal cortical neurons (*15, 16, 54*). Representations of confidence in motor thalamus were further delayed compared to pallidum and STN, beginning 40 ms prior to movement onset. This timing implies that confidence representation in the input nodes of pallidothalamic circuit is inherited from upstream cortical or striatal regions and propagated through the circuit to motor thalamus. Confidence encoding remained significant throughout the entire movement epoch and persisted into the post-feedback period in all areas (p<0.05 across 100% of the movement period; Fig. 3A,F).

In addition to systematically increasing confidence encoding latency across the pallidothalamic circuit, the temporal stability of confidence representation varied across nodes, increasing progressively from STN to motor thalamus. In STN, the encoding axis for confidence during early movement poorly predicted confidence later in the movement, and vice versa, indicating a dynamic population code. However, these changes unfolded over a relatively slow timescale (time constant for the prediction accuracy after movement onset, 580 ms; Fig. 3H,J and Fig. S5). Temporal stability of the code increased in GPi (time constant, 640 ms) and peaked in motor thalamus (time constant >1000 ms). This stable confidence code in motor thalamus extended to multiple epochs: a single encoding axis reliably decoded confidence throughout both movement and feedback epochs, with minimal variation between decoders trained during the movement and 500 ms post-feedback (Figs. 3J and S5). This progression toward a temporally stable, low-dimensional confidence code supports sequential computation and consolidation of confidence signals along the pallidothalamic circuit.

A similar gradient was observed for movement-related signals. We focused on two components: movement direction— captured by the choice encoding axis—and the transition from pre-movement to movement—the baseline modulation axis defined by the vector of intercept coefficients across neurons. As explained earlier, average neural activity rose around movement onset in all areas. This led to the reversal of the baseline modulation axis between the stimulus-viewing and movement epochs (Fig. 3C). In GPe, this transition began well before movement onset (start, 660 ms before movement onset) and ramped gradually over 1140 ms to its maximum. In subsequent nodes, the transition systematically shifted to later onsets and became faster, culminating in motor thalamus, where it started 150 ms before movement onset and ramped to maximum within only 330 ms (Fig. 3C,I). The axes defined at different times of the movement and feedback epochs were strongly aligned with each other, and anti-correlated with the axes defined during stimulus-viewing (Fig. 3C).

This provided a stable, low-dimensional signal for downstream targets of motor thalamus indicating when the movement is ongoing.

Choice was also encoded in all areas albeit more weakly than confidence, and transiently in GPe and GPi, as shown in Fig. 3B,E,F (average decoding accuracy during movement period: GPe, 55%; GPi, 54%; fraction of movement period with significant decoding: 54% in both). In contrast, STN and motor thalamus exhibited robust and persistent choice signals (average decoding accuracy: 67% in both; fraction of movement period with significant decoding: STN, 87%; motor thalamus, 93%). Further, in motor thalamus, the choice axis was stable (time constant: 500 ms, Fig. 3H and K and S5), whereas in upstream STN, it changed rapidly (time constant: 210 ms).

To determine how cognitive and motor signals became linked, we used the choice and baseline modulation axes to decode confidence and vice versa (cross-variable decoding; Fig. S6). In GPi, the baseline modulation axis, which signaled whether movement was ongoing, also decoded confidence during the movement period (baseline modulation axis decoding accuracy: 74% vs confidence axis: 77%), indicating that the two axes were well aligned. In motor thalamus, decoding confidence along either the choice axis or the confidence axis yielded the same accuracy (67%), again suggesting alignment of the two axes. GPe showed an intermediate level of alignment between cognitive and motor codes (confidence decoding accuracies: confidence axis, 69%; choice axis, 59%; baseline modulation axis, 62%). In contrast, in the STN, neither the choice nor baseline modulation axes predicted confidence (decoding accuracies: confidence axis, 72%; choice axis, 51%; baseline modulation axis, 55%), reflecting independent confidence and movement codes.

The progressive alignment of confidence and movement codes from STN to GPi and ultimately motor thalamus indicates the emergence of a lower-dimensional representation in which choice and confidence are integrated in an orderly format (see Supplementary Text and Fig. S7) and jointly broadcast to downstream regions. This finding supports the hypothesis that the pallidothalamic circuit binds cognitive variables to ongoing actions for coherent behavioral control.

### Persistent encoding of confidence and feedback after movement

Keeping track of ongoing actions, their underlying reasons, and their expected outcomes is essential for interpreting actual outcomes for strategizing and guiding future behavior (*55, 56*). Consistent with this role, confidence (i.e., the subjective likelihood of a positive outcome) and choice were stably encoded at feedback and during the following one-second intertrial interval. All nodes of the pallidothalamic circuit encoded confidence (fraction of inter-trial interval with significant decoding: GPe, 95%; STN, 100%; GPi, 100%; motor thalamus, 100%), while choice could be continuously decoded in STN and motor thalamus (fraction of inter-trial interval: STN, 84%; motor thalamus, 100%).

In STN and motor thalamus, the confidence and choice axes showed minimal dynamics in the period leading up to feedback (time constants >900 ms, Fig. S5), indicating stable codes for both confidence and its associated, recently-completed action. Thus, in STN and motor thalamus, the neural codes for movements and their expected outcomes stabilized and persisted beyond movement completion, suggesting neural circuitry primed to integrate the actual outcomes of those movements. Indeed, significant fractions of neurons in STN (17%) and motor thalamus (35%) modulated their firing rates with trial outcome (correct vs. incorrect, p<0.001) after subjects received feedback (Fig. 2C). In contrast, few neurons in GPe (6%, p=0.24) and GPi (0%, p=0.45) reflected the feedback.

Overall, pallidothalamic neurons not only show persistently elevated firing rates during movement, but also encode the confidence of *ongoing* actions, confidence in recently *completed* actions, and the *outcomes* of those actions—linking prospective evaluation with retrospective assessment.

### Emergence of reward prediction error signals in the pallidothalamic circuit

A foundational principle of learning theory is that reward prediction errors (RPEs)—the mismatch between expected and actual outcomes—drive behavioral adaptation (*8*). Expected outcomes are reflected in confidence, since higher confidence predicts correct choices and positive outcomes (*15, 16*). Thus, the representation of both the expected and the actual outcomes of decisions by pallidothalamic neurons (Fig. 4) raises the possibility that this circuit contributes to the computation of RPEs.

**Figure 4:**
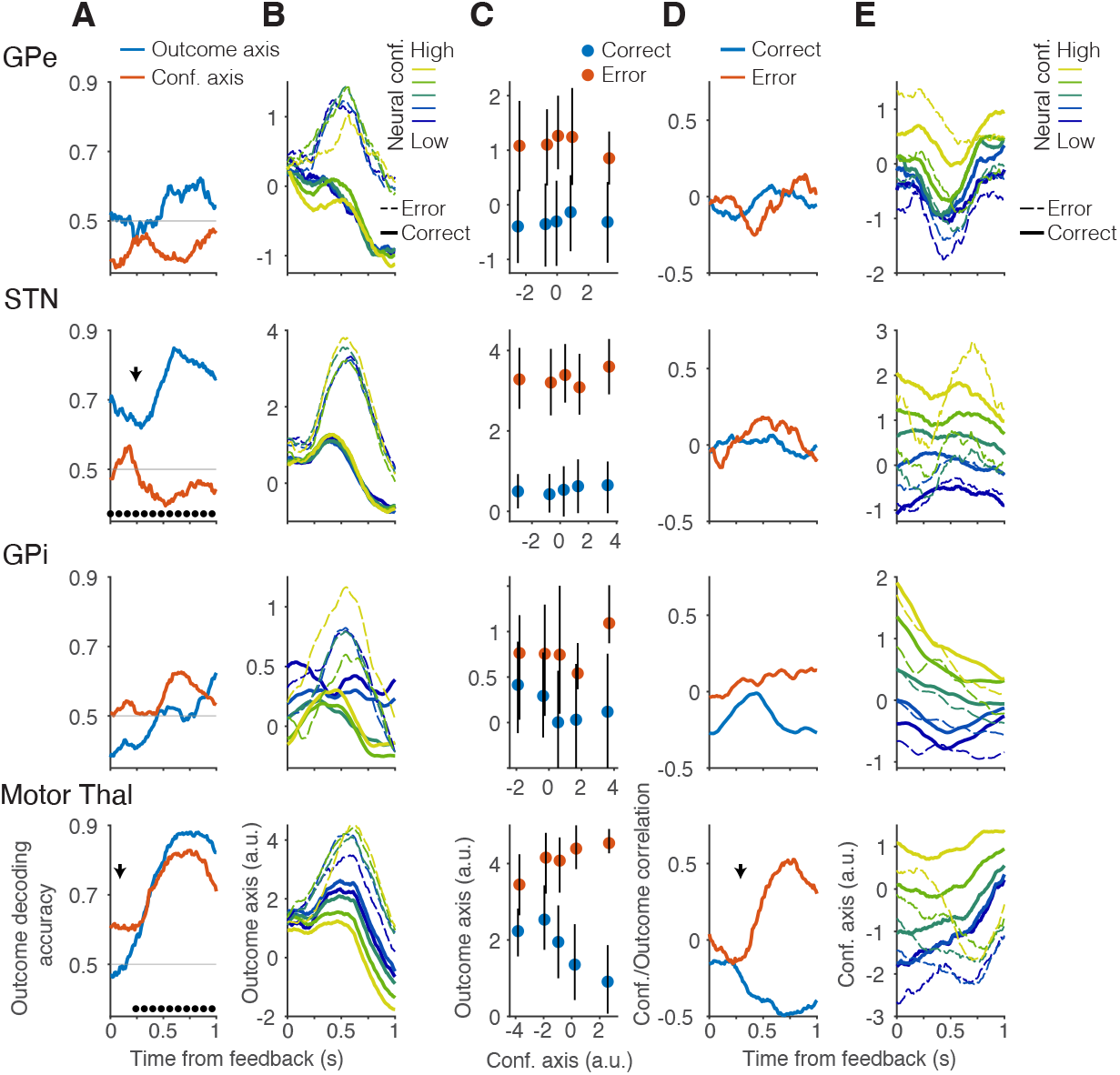
Integration of confidence and outcome into unsigned prediction error in motor thalamus. (**A**) Trial outcome encoding after feedback in STN and motor thalamus (blue). Arrows and asterisks indicate the onset and duration of above-chance decoding, respectively. In motor thalamus, trial outcome could also be decoded using the post-feedback (red) confidence-encoding axes. (**B**-**C**) Projections along the outcome-encoding axes (defined from firing rates 600 ms after feedback) for different outcomes and neurally-inferred confidence levels. The outcome axis is orthogonalized relative to the pre-feedback confidence axis (400 ms before feedback), which was used to define neurally-inferred confidence levels. In motor thalamus, confidence levels separated along the outcome axis, indicating linked representations of expected and realized outcomes. (**D**) In motor thalamus, correlations between pre-feedback confidence and outcome representations reversed over time, consistent with the representation of unsigned reward prediction errors. This transformation began 290 ms after feedback (arrow), and was absent in pallidum and STN. (**E**) Across the pallidothalamic circuit, the pre-feedback confidence axis maintained a representation of expected outcome after feedback. In motor thalamus, this axis additionally differentiated correct and error outcomes.

As described above, confidence signals persisted through movement and into the feedback epoch. These expected outcome signals were present across all pallidothalamic nodes but became remarkably stable in STN and motor thalamus—the circuit’s input and output nodes—during the period leading to feedback, providing a steady representation against which actual outcomes can be compared.

Actual outcome signals emerged selectively in STN and motor thalamus after feedback delivery. They appeared rapidly (STN: 250 ms after feedback, motor thalamus: 60 ms) and persisted with high strength (>80% decoding accuracy, STN: 460-840 ms, motor thalamus: 410-940 ms after feedback; mean decoding accuracy after feedback, STN: 75%, motor thalamus: 75%; Fig. 4A). In contrast, outcome representation was largely absent in GPe and GPi (mean decoding accuracy after feedback: 54% and 50%; p>0.05), suggesting that STN and motor thalamus do not inherit these signals from the striatum or pallidum but instead receive feedback-related inputs from dopaminergic or cortical sources.

Critically, the relationship between expected and actual outcome representations differed between STN and motor thalamus. In STN, outcome and confidence-encoding axes remained largely independent throughout the inter-trial interval (outcome decoding accuracy by the outcome axis: 74%; by the post-feedback confidence axis: 46%; mean decoding accuracy during periods of significant decoding). In motor thalamus, however, these representations converged: outcome decoding was similar along the outcome axis (78%) and the post-feedback confidence axis (75%). We hypothesized that this convergence enables computation of a reward prediction error (RPE) signal.

Two forms of RPEs have been identified. Signed RPEs are calculated as the difference between actual and expected rewards. They indicate whether outcomes are better or worse than expected and drive learning by reinforcing or discouraging behaviors. Midbrain dopaminergic neurons primarily encode signed RPEs (*57, 58*), driving the representation of RPEs in striatum and influencing reinforcement learning by updating action values based on prediction errors. In contrast, unsigned RPEs encode the absolute value of the difference between expected and actual outcomes, reflecting the magnitude of surprise, regardless of valence, and correlating with arousal, attention, and adaptive adjustment in learning rates (*53, 59, 60*). Signed and unsigned RPEs can be distinguished by how they vary with confidence or expected reward. Signed RPEs decrease monotonically with confidence on both correct and error trials. Unsigned RPEs, by contrast, are highest when high-confidence choices are followed by negative feedback, and lowest when high-confidence choices are confirmed by positive feedback. To determine whether either form of RPE is represented in the pallidothalamic circuit, we examined the relationship of pre-feedback confidence signals and orthogonal post-feedback actual outcome signals, disentangling representations of expected and realized outcomes (Fig. 4B).

In STN, confidence and outcome signals remained independent. In motor thalamus, by contrast, projection of neural activity onto the outcome axis correlated with confidence signals in a pattern matching unsigned RPE: outcome signals decreased monotonically from high-confidence errors (high surprise) to low-confidence errors, and then to low- and high-confidence corrects (Fig. 4B-C). We used pre-feedback confidence signals as a proxy for expected outcome, since confidence is predictive of choice accuracy and reaction time (Fig. S8). Outcome signals were positively correlated with pre-feedback confidence on error trials and negatively correlated on correct trials. These correlations emerged 290 ms after feedback and peaked 680 ms after feedback (r=0.52 and −0.38, respectively, Fig. 4D). Thus, motor thalamus first represented actual outcomes independently of confidence, then gradually transformed this code into one consistent with unsigned RPEs.

Conversely, the motor thalamus confidence code transformed after feedback, mixing representations of expected and actual outcomes. On correct trials, confidence-encoding firing rate modulations are stably maintained after feedback. In contrast, on error trials, movement-period activity along the confidence axis gradually transitions to the location of low confidence trials, suggesting an update to the neural representation of the expected outcome (Fig. 4E). This does not occur in STN, which throughout the inter-trial interval maintains a stable representation of confidence regardless of outcome. Thus, after feedback, there is a dichotomy between independent representations of expected and actual outcomes in STN and their gradual integration into RPE in motor thalamus.

We used partial least squares (PLS) regression to determine if an alternative linear combination of confidence and outcome could better explain motor thalamus firing rates. Before feedback, the top PLS dimension was dominated by confidence. After feedback, however, unsigned reward prediction error, which cannot be expressed as a linear combination of outcome and confidence, became the dominant component of the top PLS dimension (Fig. S9). The variance of firing rates captured by this dimension increased progressively throughout the post-feedback window, peaking 710 ms after feedback, consistent with a gradual emergence of an unsigned RPE signal. Thus, in motor thalamus, the output of the pallidothalamic circuit, firing rates evolved to encode an integrated quantity nonlinearly constructed from confidence and outcome.

STN and motor thalamus maintain distinct representations of action outcomes. In motor thalamus, actual and expected outcomes are integrated into unsigned RPEs. In STN, actual and expected outcomes are separately maintained. In contrast, GPe and GPi had minimal representation of the realized outcome and were devoid of RPE signals, suggesting that the unsigned RPE signals in motor thalamus are not transformations of the signed RPE representation in the striatum. Instead, they likely emerge within motor thalamus via convergent integration of confidence and outcome signals.

## Discussion

Traditional views of the basal ganglia and motor thalamus emphasize a feedforward architecture in which the pallidothalamic pathway gates motor commands, relaying selected actions to cortex while suppressing competing actions. Yet, real-world behavior requires more than motor gating—it demands continuous coordination between movement and cognition.

In the first neuronal recordings across the human pallidothalamic circuit during decision-making and movement, our results challenge this prevailing framework and reveal the circuit as a dynamic and integrative hub, where neurons persistently encode not only motor execution but also cognitive variables associated with the movement, including confidence, reward outcome, and prediction errors. The representation of cognitive signals was neither transient nor stereotyped. Rather population-level codes for movement, choice, and confidence were robust, low-dimensional, and stable, particularly in the motor thalamus. Across the circuit, latency, stability, and alignment of motor and cognitive codes increased from GPe and STN to GPi and motor thalamus, consistent with a hierarchical organization in which early-stage nodes contribute dynamic computations that are consolidated into persistent representations in later stages. This hierarchical transformation was especially striking for confidence and outcome signals. Confidence emerged during deliberation and persisted through movement and feedback. The representation of confidence, however, emerged only in the late stimulus-viewing period, suggesting that the pallidothalamic circuit does not compute confidence directly but instead receives it from upstream cortical and striatal regions, and binds it to the relevant action plan. Embedding confidence within movement-related population dynamics also enables online control policies that adjust commitment and vigor during execution (*49, 55, 61–63*).

In contrast, outcome encoding arose shortly after feedback but was largely absent in pallidum. As a result, unsigned reward prediction error (RPE)—the mismatch between expected and realized outcomes—emerged in motor thalamus, but not in GPe or GPi. The lack of outcome and RPE signals in pallidum suggests that these computations are unlikely to have been inherited from striatum but rather arise locally or via independent afferents. More broadly, maintaining confidence and outcome signals through movement and juxtaposing them with feedback provides a substrate for temporal credit assignment—linking outcomes to the specific cognitive-motor episode that produced them (*7–9*).

We propose that this integration of cognitive and motor signals in the pallidothalamic circuit enables a critical but underappreciated computation: supporting “acting while deciding” and “deciding while acting.” Stable confidence and outcome codes during and after movement, and their convergence into unsigned RPE, point to a system evolved not merely for gating but for maintaining and updating internal models of action-confidence and action-outcome contingencies. Everyday behavior illustrates this need: we can walk and simultaneously converse, while adjusting pace to avoid obstacles and refining our arguments. Such abilities require binding cognitive elements to their corresponding actions, preventing interference across tasks. Whereas the perceptual binding problem—linking features like color and shape into a unified object—has long preoccupied neuroscience, the analogous cognition–action binding problem has received far less attention. Our results suggest that even in the output nodes of the basal ganglia–thalamus circuit, traditionally regarded as motor-specific, cognitive variables are integrated with actions to resolve the binding problem.

An enticing possibility is that the anatomical position of motor thalamus—at the interface of basal ganglia and cortex— enables it to act as a key broadcast node for cognitive-motor integration. With widespread projections to motor cortex and other frontal areas (*64*), motor thalamus is well-positioned to distribute information about action timing, confidence, outcome, and prediction errors, coordinating computations across frontoparietal networks and shaping cortical responses based on recent behavioral outcomes. The presence of unsigned RPE in motor thalamus suggests that higher-order thalamic nuclei may serve as a critical source of RPE signals in cortex, informing distributed learning and adaptation. These functions could be facilitated by extensive inputs to these thalamic nuclei (*64,65*) as well as convergent sensorimotor and limbic cortico-basal ganglia loops (*66*).

Together, these findings advance a new view of the pallidothalamic circuit as a substrate for hierarchical, persistent, and integrated representations that bind motor actions to their cognitive variables. By challenging the classical view of basal ganglia and motor thalamus as mere gating mechanisms, our results highlight their broader role in the interplay between decision-making and motor execution that characterizes flexible behavior.

## Methods

All study procedures were approved by the NYU Langone Health Institutional Review Board. Subjects were patients with Parkinson’s disease (PD, 8 patients) or essential tremor (ET, 4 patients) undergoing implantation of deep brain stimulation (DBS) electrodes. Patients were evaluated pre-operatively in our outpatient clinic by a multidisciplinary team—including movement disorder neurologists (LP, MP), a neurosurgeon (AYM), and a neuropsychologist—following standard clinical protocols. Only patients with no more than minimal cognitive impairment who expressed strong interest in participating were enrolled after providing informed consent. Subjects completed training on the behavioral tasks in the weeks preceding surgery, achieving stable behavioral performance and comfort with the paradigms. Surgical procedures followed our standard clinical protocol (*48*), with patients awake and off their usual dopaminergic or tremor-suppressive medication. Neural recording sites were localized using intra-operative computed tomography (CT) co-registered to pre-operative structural MRI, along with microdrive positioning data and intra-operative neurophysiological features verified by skilled movement disorder neurologists (LP, MP).

While prior human studies have investigated individual nodes in the pallidothalamic circuit, here we sampled the motor thalamus together with its pallidal and subthalamic afferent nodes. Interrogating these nodes within the same experimental setup is crucial to understanding circuit-level computations, but requires a diverse patient population. We use several strategies to account for the heterogeneity of diseases (PD vs. ET), symptoms in patients with different DBS targets (STN vs. pallidal targets in PD), and demographic and clinical factors.

We previously characterized normal human performance on our main task (*67*). Trained patients exhibited psychometric, chronometric, and confidence functions that (1) varied systematically with stimulus strength, (2) were similar across diseases and targeted regions, and (3) replicated trends observed in neurotypical individuals.

Our main findings were consistent across disease types and DBS targets, establishing that our results reflect the properties pallidothalamic circuit not the disease profile. In all patients, movement was associated with elevated firing rates and in 11/12 patients, neural activity was modulated by both cognitive and motor features. After movement, neural activity in STN and motor thalamus robustly represented confidence and feedback. Moreover, systematic differences across nodes (e.g., neural latencies of movement and cognitive signals) were consistent with known anatomical connectivity. Replication of these features across patients with different DBS targets, diseases, and symptom profiles supports their generality as properties of the human pallidothalamic circuit.

### Task design

Subjects performed visual discrimination and reaching tasks during micro-electrode recordings conducted for clinical localization of DBS targets. The visual discrimination tasks required subjects to categorize either facial expressions (e.g., happy vs. sad) (*52*) or the net direction of motion (rightward vs. leftward) of a random dot stimulus. Subjects jointly reported their choice and confidence (*50, 67*) with a single reaching movement. Hand position was recorded in three dimensions at 110 Hz with sub-millimeter precision (*68*) using a stereoscopic infrared camera (Leap Motion controller). This camera controlled cursor position on the stimulus display, which was mounted on an adjustable arm to give the patient an unobstructed view without interfering with the surgical field.

Each trial began when the subject moved their hand to bring the cursor into a central fixation area indicated by a white circle. Four targets then appeared, one in each corner of the screen. Left and right targets corresponded to the two possible categories (e.g., happy or sad), while target colors on each side indicated confidence level (green, high confidence; red, low confidence). After a variable delay (0-0.5 s, sampled from a truncated exponential distribution), the face or random-dot stimulus appeared above the fixation area and stayed on until response initiation. Subjects indicated both their choice and confidence about the choice with a single reach to one of the four targets. We used binarized choices and confidence reports mapped to well-separated target locations to ensure reliable reporting even in patient with tremor.

Subjects were instructed to maintain gaze fixation during stimulus presentation and to initiate their response as soon as they were ready. The stimulus was extinguished when the hand left the fixation area or earlier if its velocity exceeded a preset threshold. Subjects received audiovisual feedback, indicating whether their choice was correct, regardless of reported confidence. Feedback onset occurred after target acquisition with a minimum delay of 400 ms and an additional delay for response times (RT) under 1000 ms (1 s− RT). Response time was defined as the interval between stimulus onset and movement initiation. Arm movement durations spanned approximately 1-2 s (median, 1.4 s; interquartile range, 1.3-1.8 s), creating a substantial separation (∼2 seconds) between decision-related processes and feedback.

We used a template-fitting approach to find where subjects’ acceleration was discontinuous, representing the point of movement onset. This template consisted of constant followed by linearly increasing acceleration. On a trial-by-trial basis, we fit the time window matched to this template and the relative duration of the constant and linear terms. A penalty term was used to prevent large variations in the template.

We computing movement onset in each trial by fitting a bilinear function to hand acceleration:

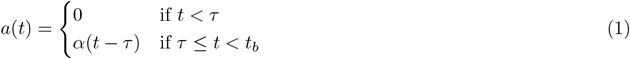

where α is the acceleration rate of increase, t_*b*_ is the right edge of the time window used for analysis, and τ is the movement onset. Acceleration, a(t), was computed as the second derivative of 3D hand position, with a smoothing step after each derivative (causal, half-Gaussian kernel, SD=91 ms), and model the best model parameters (τ, t_*b*_, α) were estimated for each trial. Detected movement onsets were visually inspected relative to acceleration traces and manually corrected if necessary. Manual corrections were performed blind to electrophysiological data and trial condition.

The stimulus was either a random-dot kinematogram (*12, 20*), or a stochastically varying face (*51, 52, 69*). Face stimuli were morphed between two prototype expressions. To reduce stimulus dimensionality, morphing was restricted to three features (eyes, nose, mouth) (*69*). On each trial, we choose a nominal morph level between the two prototypes (−96% to +96%). This nominal morph level determined mean morph of the three informative features and the correct choice. Features values fluctuated independently every 106.7 ms according to a Gaussian distribution (SD, 10% morph). Feature changes were masked using a sample-mask cycle: stimulus shown for 13.3 ms and then gradually faded out as a mask with similar spatial frequency content faded in. In the next cycle, a new face stimulus with slightly altered informative features was shown, followed by fading with another mask, and so on. The stimulus in each trial looked like a face appearing and disappearing behind cloud-like noise patterns. The masking procedure was effective, making feature changes subliminal, yet the stimulus fluctuations influenced behavior. (*52, 69*).

The Reach task was randomly interleaved with the Decision trials. Each Reach trial began when the subject moved their hand into the fixation area, followed by the appearance of a *single* target in one corner of the four screen corners corresponding to Decision-task target locations. Subjects then reached to the target and received audiovisual feedback.

### Electrophysiological Recordings

Micro-electrodes (Neuroprobe and Alphaprobe Sonus Shielded tungsten microelectrodes; Alpha Omega, Alpharetta, GA) consisted of multiple recording surfaces: the sharp electrode tip (250-1250 kΩ impedance) capable of recording sortable single units, four round micro-contacts radially spaced around the electrode 2.5-2.75 mm above the tip (70-800 impedance kΩ, Alphaprobe only), and macro-contacts located 3 mm and 4.5 mm above the tip (1-10 kΩ impedance).

According to our standard clinical protocol, DBS electrodes were implanted in STN or GPi in patients with Parkinson’s disease and in the Vim subregion of the motor thalamus (*70*) in patients with essential tremor. Micro-electrode recordings were performed at multiple locations along a linear trajectory including within the GPe (during GPi-targeted surgeries). Overall, we recorded from neurons in STN (5 hemispheres, 50 neurons), motor thalamus (5 hemispheres, 46 neurons), GPi (4 hemispheres, 12 neurons), and GPe (5 hemispheres, 34 neurons).

### Spiking Data Analysis

Spike detection and sorting were performed with an offline sorter (Plexon, Dallas, TX) using raw electrophysiological recordings from the micro-electrode tip contact sampled at 44 kHz. Spike sorting was performed manually using principal components of spike waveforms. Single units, multi-units, and unsorted background spiking were treated as separate units. Single units were defined as those with the clearest isolation from background and most consistent action potential waveforms (29 units). Spike-counts in 10 ms bins were convolved with a causal kernel (alpha function, time constant 200 ms) to estimate instantaneous firing rate.

Recording surfaces along the electrode shaft captured background spiking activity that could not be sorted into single units. We analyzed these signals using entire spiking activity (ESA) (*71*), a threshold-free measure of firing rate. Raw voltage sampled at 44 kHz was filtered forward and backward using a 16th-order 300 Hz high-pass Butterworth filter and full-wave rectified. The result was convolved with the causal alpha kernel and averaged in 10 ms bins to calculate instantaneous firing rates.

Thus, our dataset included single-unit, multi-unit, and background spiking activity. Analyses performed separately across these data classes yielded comparable results and were, therefore, combined. For simplicity, unless otherwise specified, we refer to all of them as units.

Instantaneous firing rates were aligned to stimulus, movement, and feedback onset. To combine data across units, firing rates were z-scored using the mean and standard deviation calculated across all responded, non-Reach trials. Analysis windows were chosen such that each time point contained data from at least 50% of trials across all trial conditions and subjects.

### Single unit analyses

To determine whether firing rates changed around movement onset, we calculated peristimulus time histograms (PSTHs) aligned to movement onset and compared mean firing rates 250 ms before versus 250 ms after movement onset relative to baseline activity measured in a 350 ms window before stimulus onset. Statistical significant was assessed using the Wilcoxon rank-sum test. To test whether changes were sustained, we divided the activity between 250 ms before movement onset and 10 ms before feedback into non-overlapping 100 ms bins and compared to baseline using Holm-Bonferroni corrected Wilcoxon rank-sum test. Neurons were classified as having sustained increases or decreases if activity significantly differed from baseline for >75% of the movement period.

Unsupervised clustering of condition-independent neural responses was performed using principal component analysis on concatenated PSTHs aligned to stimulus onset, movement onset, and feedback. Ward’s method was applied to loadings from the top principal components representing >85% of the variance.

We used a cluster-mass permutation test (*48, 72*) to identify when single-unit firing rates encoded cognitive variables (choice, confidence, outcome), controlling for multiple comparisons. We isolated the effects of specific cognitive factors or their interaction by performing two-factor cluster-mass tests with pairs of variables (confidence/choice, confidence/outcome) on single-trial firing rates aligned to stimulus, movement, or feedback onset. Null distributions were generated by permutation of residuals under reduced models (*73*). Effect sizes were computed by averaging firing rates across significant time points and then across trials.

We compared the fraction of neurons in each area encoding a task variable within one or more *a priori* time windows to the expected fraction under a binomial distribution (p=0.05). These windows included the stimulus period, extending for half the median reaction time after stimulus onset, the movement preparation time, extending for half the median reaction time before movement onset, the movement period, extending from movement onset to feedback, and the one-second post-feedback period.

Choice-confidence interaction effects were further examined using the Wilcoxon rank-sum test on single-trial firing rates. Four comparisons were performed: high vs. low confidence on contralateral trials, high vs. low confidence on ipsilateral trials, contralateral vs. ipsilateral low confidence trials, and contralateral vs. ipsilateral high confidence trials. Single-trial firing rates were calculated across time points identified by the cluster-mass test and compared with Holm-Bonferroni corrected rank-sum tests. Neurons were classified as selective for confidence, choice, or both (p<0.05).

Time windows exhibiting significant confidence- or choice-related firing rate modulations on decision trials were compared to analogous windows on Reach trials. Reach and Decision effect sizes were compared within neurons using the Wilcoxon signed-rank test.

To determine whether confidence-related firing rate modulations could be explained by movement kinematics, we sampled trials to equalize the distribution of hand velocity across high- and low-confidence conditions. Sampling probabilities for pairs of high- and low-confidence trials were chosen to minimized their velocity difference:

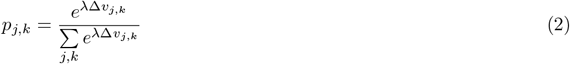

where 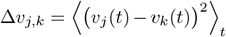 and λ was set to 10. We selected 100 trial pairs per session and quantified confidence-related firing rate modulation together with residual speed differences within time windows exhibiting significant confidence effects (Fig. S4).

### Neural population analyses

To quantify the strength, stability, and dynamics of population codes for task variables, we constructed linear decoders, 𝒟_*x*_, that classify trial-by-trial neural population activity into binary classes of a variable x ∈ {−1, 1}, representing choices (right/left), confidence (high/low), or outcome (correct/error). Let 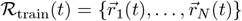 denote the population activity matrix, where 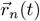 is the vector of instantaneous firing rates of neuron n across trials at time t, excluding two held-out trials. Because neurons in our dataset were recorded across sessions, we created pseudo-populations by combining trials with similar stimulus strengths, choices, confidence levels, and outcomes across sessions.

To identify decoding axes in the population state space, we fit a linear encoding model to each neuron’s firing rate modulations in the training set:

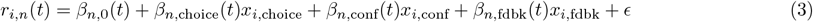

where x_*i*_ represent task variables (choice, confidence, or feedback) on trial i, and β_*n*_(t) are model coefficients. We concatenated coefficients across neurons to derive encoding axes, which define linear discriminant directions:

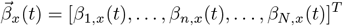

The projection of population activity patterns along these axes forms a linear discriminant, q:

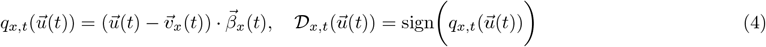

where 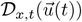 denotes the decoded task variable from population activity at time t, 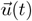is the recorded activity, and 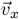 is the population activity predicted by a reduced encoding model excluding the target task task variable x. Put differently, we project the residual firing rate modulation unique to the target task variable onto the encoding axis derived from the training data.

To assess decoder performance, we held out two trials to form the test set,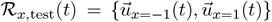 where 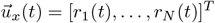 and each *r*_*n*_(*t*) is the instantaneous firing rate of neuron *n* on a single trial within the matched pseudo-population. The linear discriminant was applied to these held-out trials. We report the decoder performance as average accuracy over cross-validation folds (500 iterations).

We evaluated *cross-temporal* decoding performance by decoding ℛ_*x*,test_(*t*) using decoders trained on different times, *τ*. We did not orthogonalize the encoding axes so that we could perform *across-variable* decoding by using the axis for one task variable to classify another variable. Cross-temporal decoding measures functional alignment of the encoding axis for the same task variable across time, whereas cross-variable decoding measures the alignment of encoding axes for different task variables.

Cross-variable decoding lacks unique ground-truth labels, as there is no objectively correct mapping between ipsilateral versus contralateral choices low versus high confidence, and positive versus negative feedback. We defined a convention to match labels for ipsilateral choice, high confidence, and correct feedback. Higher-than-chance cross-variable decoding accuracy therefore indicates that ipsilateral choices, high confidence, and positive feedback modulate firing rates in the same direction in state space, whereas lower-than-chance accuracy indicates modulations in opposite directions. Near-chance accuracy indicates that encoding axes are approximately orthogonal.

To characterize firing rate dynamics independent of cognitive variables (Fig. 3C), we constructed baseline axes using the intercept coefficients from Eq. 3. These axes capture condition-independent rate changes related to presence or absence of movement. To avoid contamination by firing rate autocorrelations, we split trials from each sessions into independent halves, fit the encoding model separately to each half, and formed baseline axes by concatenating intercept coefficients β_0,*n*_ across neurons. We the computed the cosine similarity between baseline axes of different times in one half and the movement-period axis from the other half. The cosine similarity was averaged across cross-validation folds (500 iterations)

To visualize neural activity along task-relevant dimensions in neural state space, we orthogonalized encoding axes for different task variables at selected time points and projected population activity onto these axes.

### Partial least squares

We used PLS regression to determine to what degree motor thalamus firing rates vary with confidence, outcome, and unsigned reward prediction error. PLS identifies low-dimensional linear combinations of firing rates that maximally covary with behavioral variables. For each trial, we defined a three-dimensional task-variable vector: 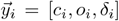, where *o*_*i*_ ∈ {1, 0} denotes correct or incorrect outcomes, *c*_*i*_ ∈ {0.6, 1} denotes low or high confidence, and *δ*_*i*_ = |*c*_*i*_−*o*_*i*_| denotes unsigned reward prediction error. Due to the binary nature of o and c, the exact numerical values assigned to them are not critical for the results. We selected 100 trials for each combination of confidence and outcome across sessions. The resulting pseudo-population firing rate vectors formed the predictor matrix X and corresponding task variable matrix Y. Columns of Y were z-scored, and both X and Y were mean-centered before computing two PLS components using MATLAB (plsregress function). PLS regression was repeated across 50 pseudo-population draws. Loadings shown in Fig S9 are the averages of normalized loadings across runs. Similarly, the fraction of variance explained was computed for each run and averaged across runs.

## Acknowledgments

We thank Bianca Sieveritz and Gouki Okazawa for help with stimulus design. We thank Cathryn Lapierre for assistance with administration and subject recruitment. We thank Anders Nelson and Bijan Pesaran for insightful discussions and feedback on the manuscript. This work was supported by the National Institute of Neurological Disorders and Stroke (R01 NS145489). R.K. was additionally supported by the Pew Innovation Fund (00037214), the National Institute of Mental Health (R01 MH127375 and R01 MH141929), and the Simons Collaboration on the Global Brain (GB-Culmination-00002986-02).

## Author Contributions

D.L. and R.K. conceptualized and designed the study and analyses. D.L., M.S., L.P., M.P., and A.M. collected the data.

D.L., A.M., and R.K. wrote the manuscript.

## Competing interests

The authors declare no competing interests.

## Data availability

All datasets generated and analyzed in this study will be deposited in a public repository and made available upon publication.

## Code availability

All analysis code will be deposited on github and made available upon publication.

## Supplementary Text

### Controlling for distinct movements

To determine if representations of confidence could be explained by neural encoding of distinct movements to high- and low-confidence targets, we asked whether analogous representations were present in the Reach task. In this task, subjects made instructed reaching movements to the same four targets as in the Decision tasks. Whereas movements to the four targets reported confidence and choice in the Decision tasks, target locations carried no such meaning in the Reach task. Although a significant fraction of neurons in all areas distinguished movements to the upper and lower targets on each side during Decision tasks (Fig. 2), this was not the case in the Reach task (fraction of neurons with significant encoding: GPe, 0% p=0.68; STN, 12%, p=0.050; GPi, 0%, p=0.30; motor thalamus, 3.9%, p=0.38). Movements to the upper and lower targets in the Reach task could not be decoded from the time windows identified as confidence-encoding in Decision tasks (fraction of confidence-encoding neurons with significant encoding of upward/downward movements: GPe, 0% p=0.40; STN, 0%, p=0.34; GPi, 14%, p=0.044; motor thalamus, 0%, p=0.37). Furthermore, confidence-associated firing rate modulations in the Decision tasks were 79-160% greater than those associated with upward and downward movements during the same time windows in the Reach task (Fig. S3).

### Controlling for movement vigor

Neurons in the pallidothalamic circuit are modulated by movement kinematics (*48, 74, 75*). Therefore, we tested whether the firing rate differences between high- and low-confidence trials could be explained by variations in movement vigor. Differences in vigor were small (3.6% difference in peak speed or 0.12 z-score units; 2.9% difference in mean speed during movement or 0.09 z-score units). Within time windows with significant confidence-related firing rate modulations, relative kinematic differences were 50-86% smaller than firing rate differences (as shown in Fig S4).

Critically, firing rate differences between high- and low-confidence trials persisted even after resampling trials to equalize their kinematic distributions across confidence levels. Notably, kinematic equalization increased the magnitude of confidence-related firing rate modulations in STN and did not significantly change it in other areas (GPe, p=0.37; STN, p=0.005; GPi, p=0.56; motor thalamus, p=0.57, Wilcoxon signed rank test).

### Cognitive-motor alignment

We visualized concurrent but dissociable firing rate modulations associated with confidence and movement to assess how cognitive and motor variables are linked in the pallidothalamic circuit. We stratified population responses by their projection onto the confidence axis and visualized their evolution along *orthogonalized* choice and baseline modulation axes (Fig. 3D and S7). Activity along these movement-related axes was concurrent with the confidence-related activity but was independent from it.

In STN, confidence and choice remained largely independent: we did not observe systematic choice-related firing rate modulations along the confidence axis or confidence-related modulations along the choice axis. Similarly, in pallidum or motor thalamus, the confidence axis did not represent the choice.

However, confidence was expressed along the choice and baseline-modulation axes in pallidum and motor thalamus (Fig. S7). In GPe and GPi, the separation of ipsilateral and contralateral trials along the choice axis was larger for high-confidence trials. The opposite was true in motor thalamus, where low-confidence trials showed the largest modulation along both the baseline-modulation and choice axes. Furthermore, on contralateral trials, confidence was represented along these movement-related axes. The emergence of such mixed representations lends additional support to the hypothesis that the pallidothalamic circuit integrates cognitive and motor codes.

## Supplementary Figures

**Figure S1:**
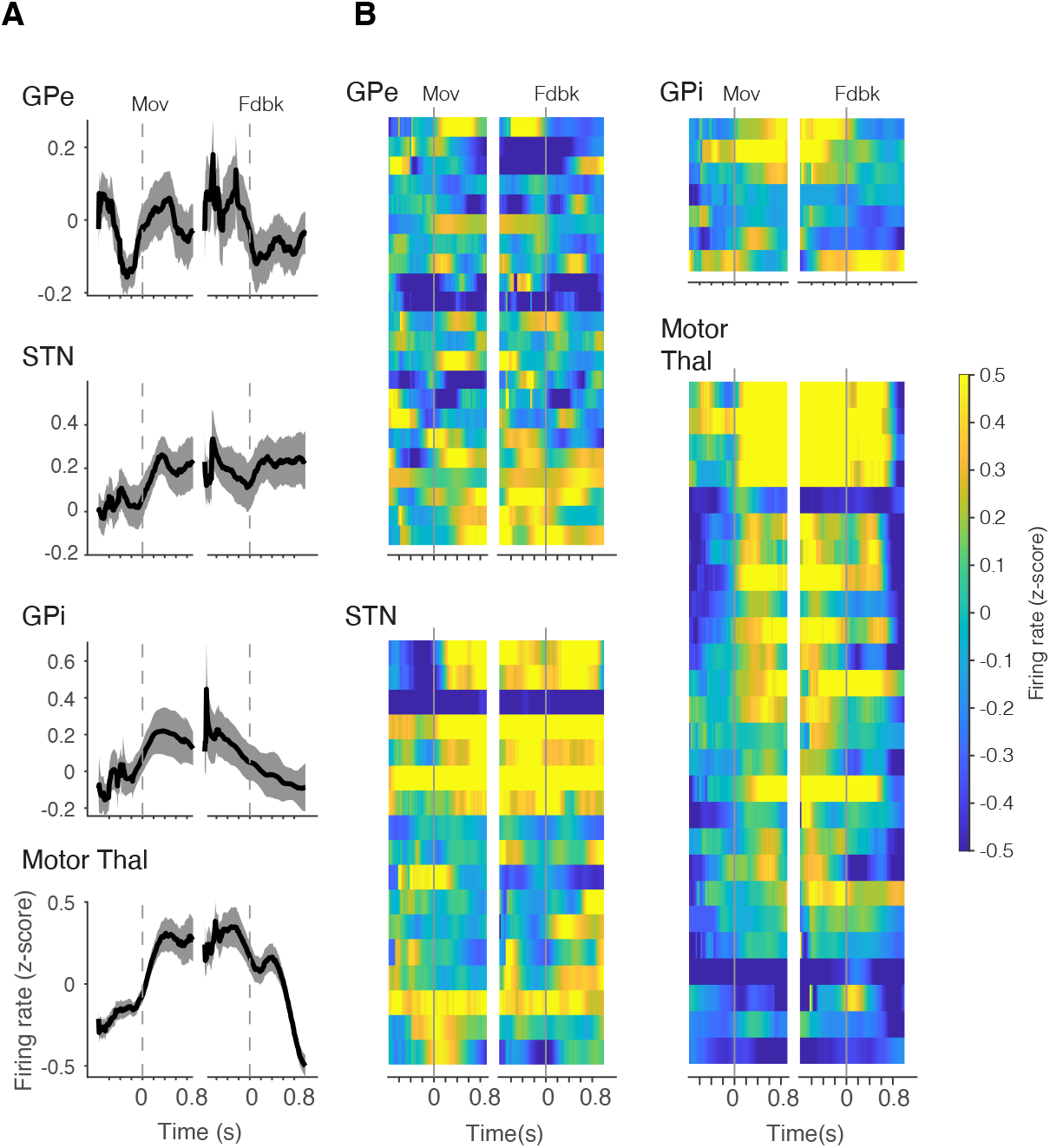
Movement-related neural activity during the instructed-reach task. (**A**) Population-averaged PSTHs in each target region. (**B**) Normalized firing rates of individual neurons aligned to movement onset and feedback.

**Figure S2:**
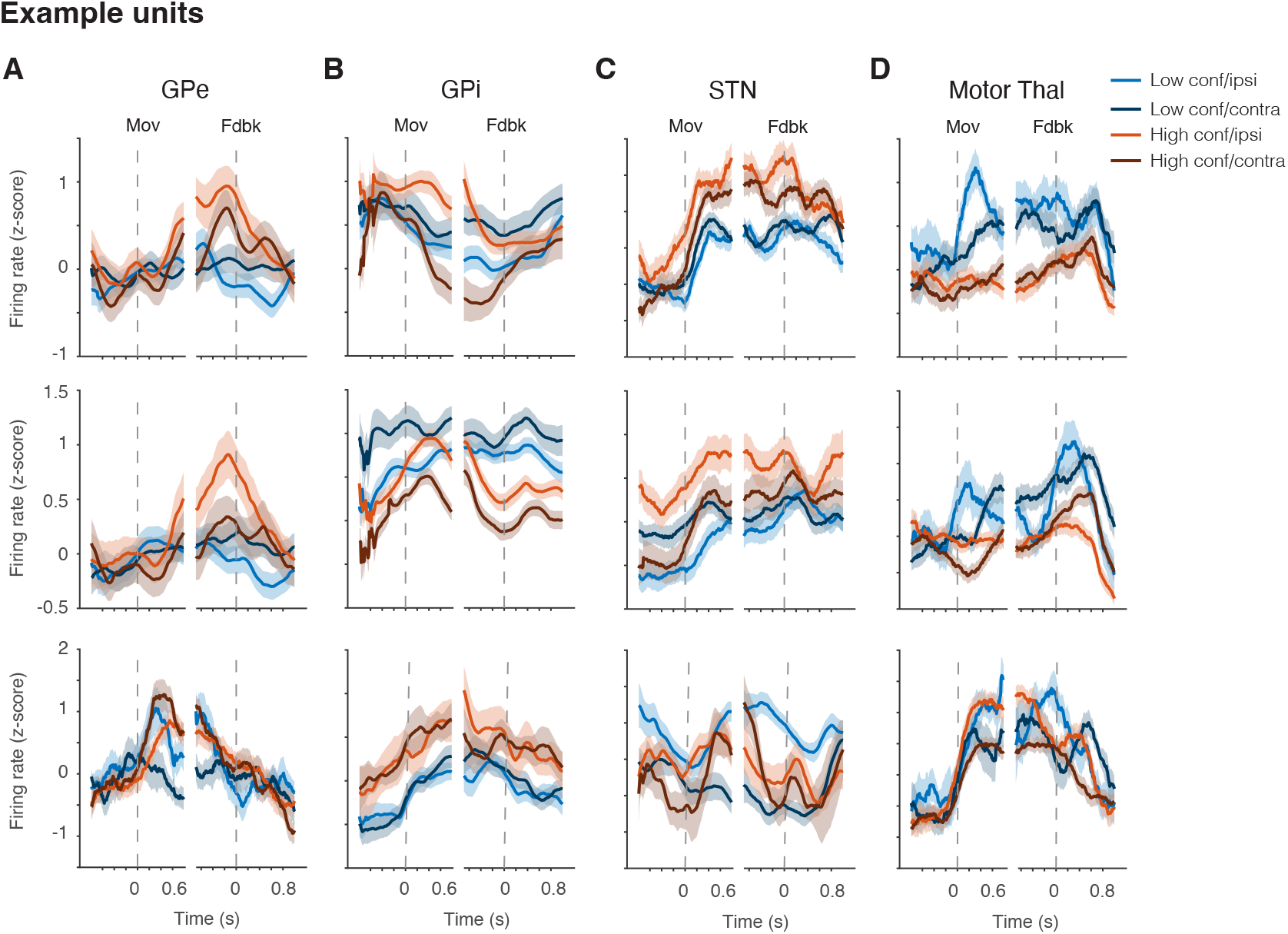
Confidence and choice are encoded by neurons with diverse firing rate response profiles. Example neurons showing firing-rate modulation by confidence or choice in GPe (A), GPi (B), STN (C), and motor thalamus (D).

**Figure S3:**
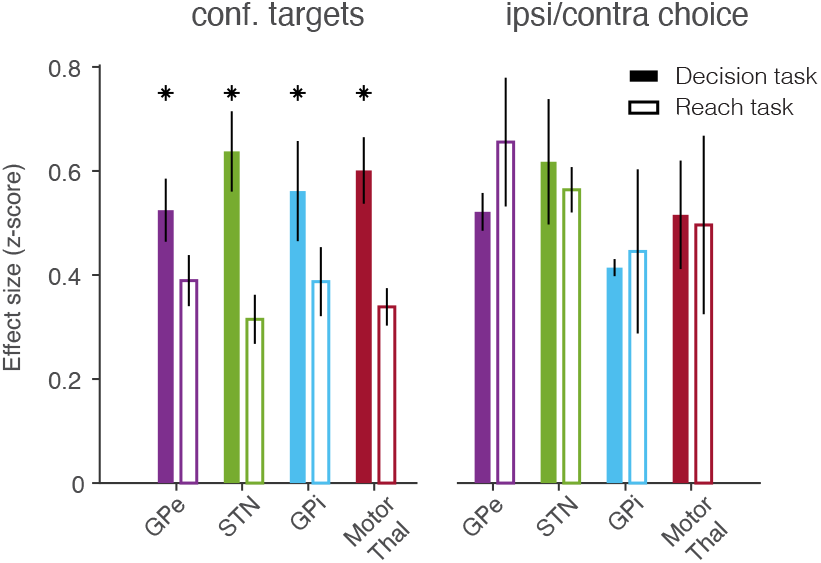
Comparable confidence- and choice-related firing rate modulations. Normalized rate modulations (z-score) during movement preparation and movement periods are shown separately for confidence and choice on decision tasks (filled bars), and for reaching to the same targets during the instructed-reach task (open bars). Only neurons recorded in both tasks are included. * indicates p<0.05, Wilcoxon signed rank test.

**Figure S4:**
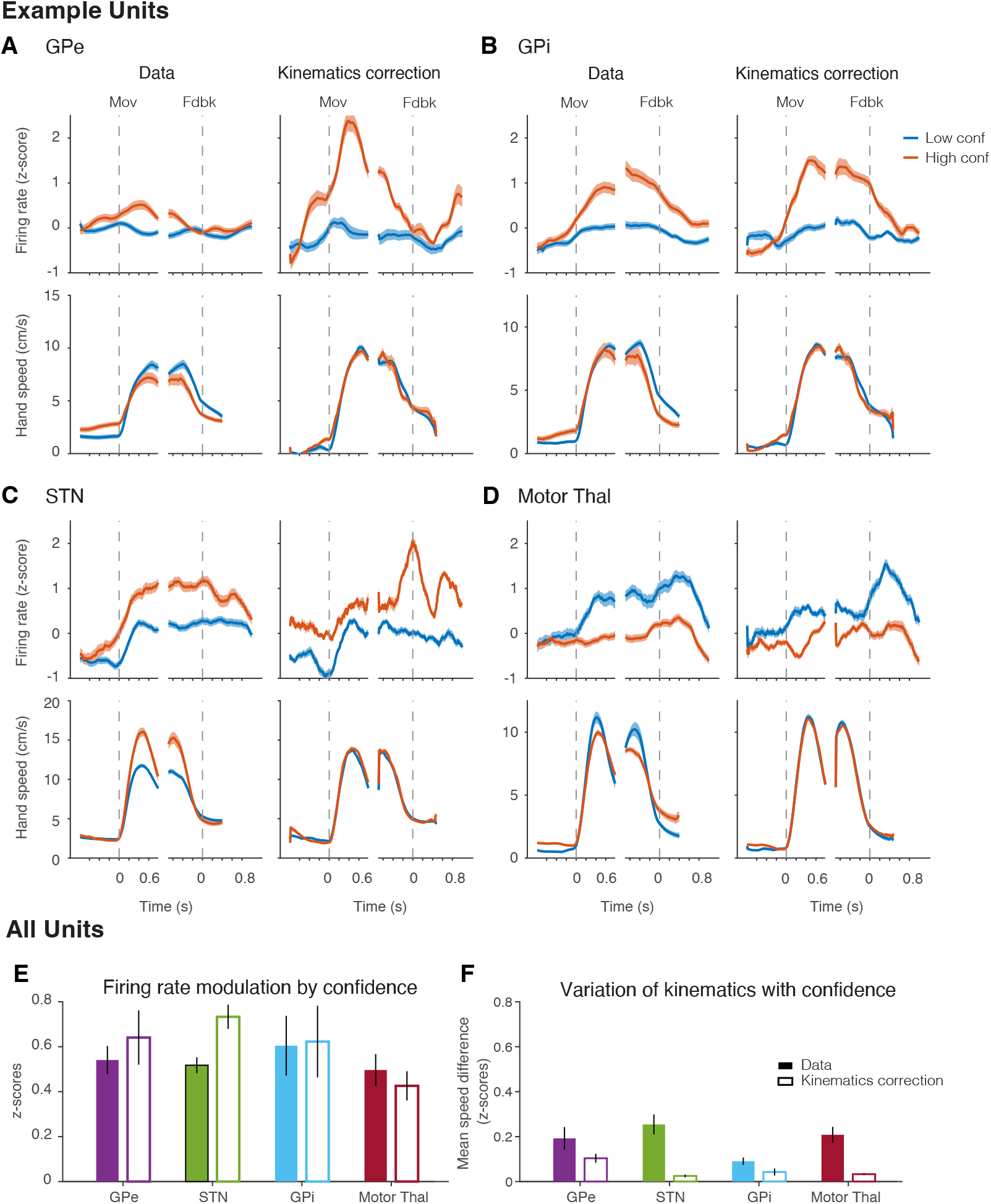
Confidence encoding is not explained by movement kinematics. (**A**-**D**) Example neurons from GPe (A), GPi (B), STN (C), and motor thalamus (D). Top: PSTHs for low- and high-confidence trials before (top left) and after (top right) resampling of trials to match movement kinematics. Bottom: corresponding hand speed profiles before and after matching hand kinematics. (**E**) Normalized confidence-related firing-rate modulation before (filled) and after (open) matching movement kinematics. (**F**) Mean difference in hand speed before and after kinematic matching.

**Figure S5:**
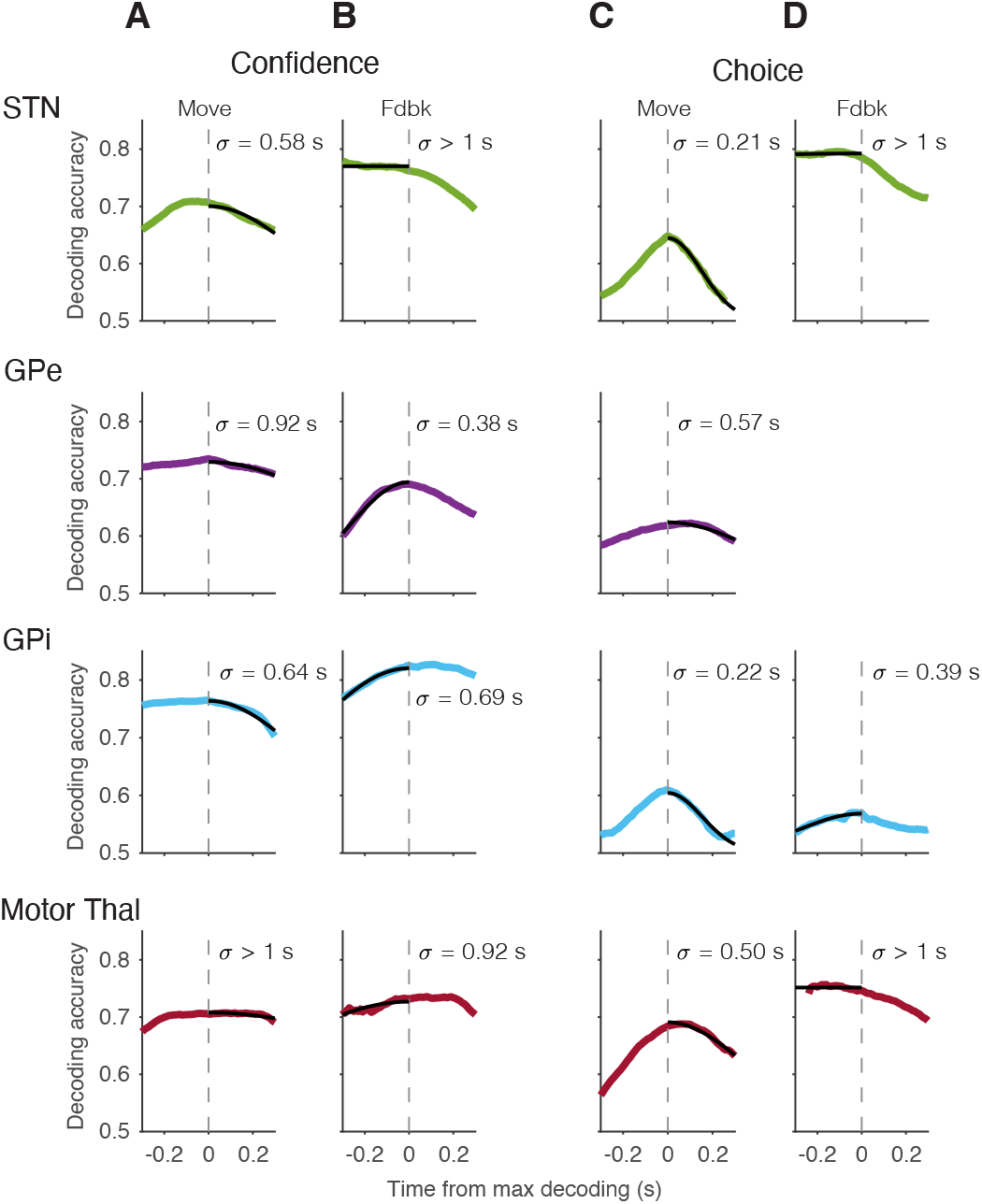
Temporal stability of confidence and choice representations. (**A**-**B**) Decoding accuracy of confidence classifiers trained on population activity at one time and tested at times before and after. (**C**-**D**) Same as A-B, but for decoding choice. Time 0 corresponds to the average decoding accuracies along the diagonals in Fig. 3E-F. Positive and negative times reflect averages of off-diagonal values with specific offsets. A and C focus on the movement window; B and D focus on a 560 ms window before feedback. Decoding accuracies represent a binary variable and were thus logit transformed prior to averaging. The inverse logit transform was applied to visualize the decline in decoding accuracy of off-diagonal decoders. Only time points with strong (above-threshold) statistically significant decoding on the diagonal were included to evaluate the dynamics of the neural code itself, rather than the strength of the code. The threshold was defined as the mean decoding accuracy of significant decoders. GPe time constant for pre-feedback choice encoding is undefined because its neurons did not reliably encode choice in this window (Fig. 3F). Time constants (σ) are the standard deviation of a Gaussian fit (black lines).

**Figure S6:**
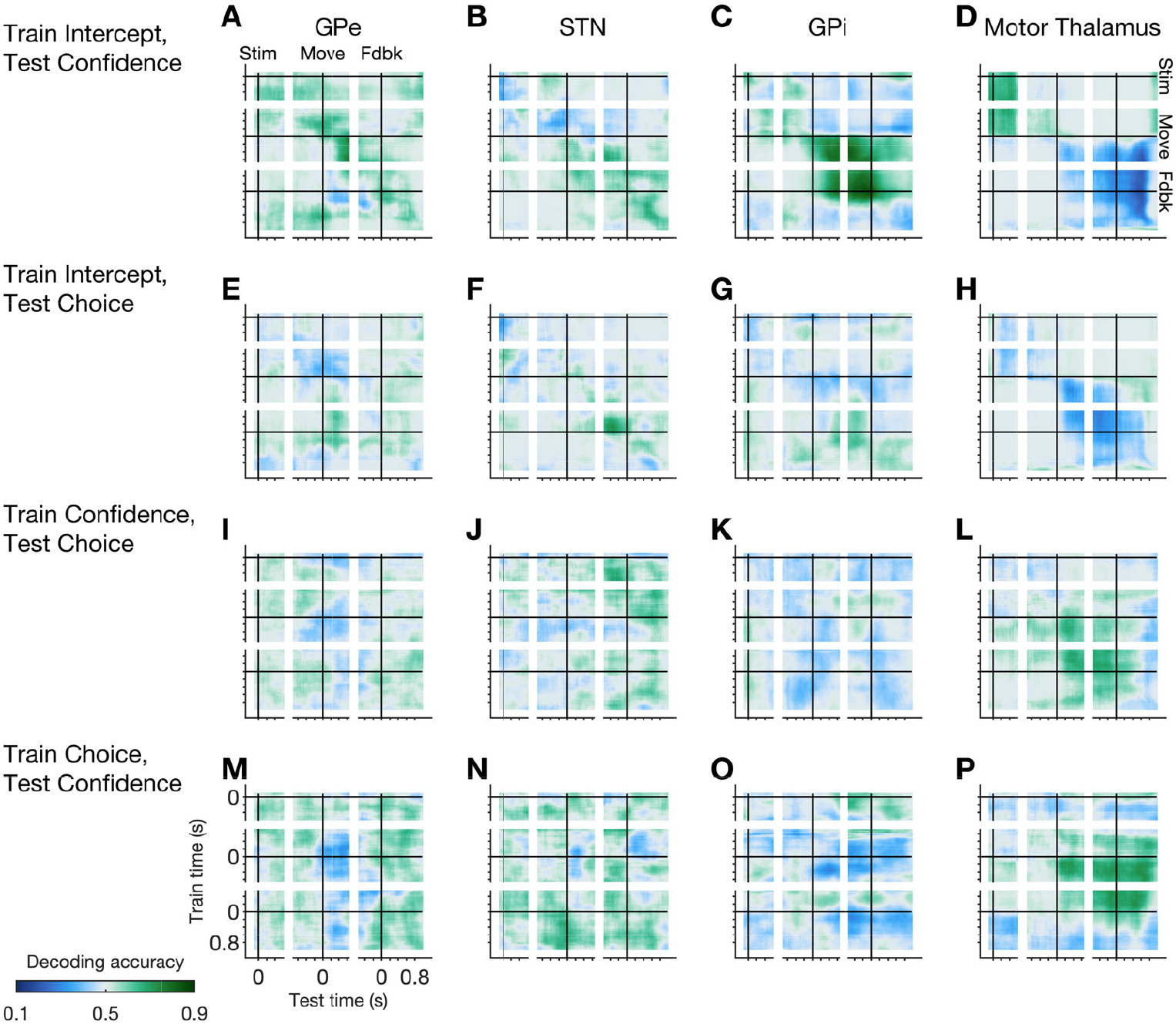
Alignment of population codes across variables. (**A**-**H**) Cross-variable decoding using axes trained on baseline modulation (intercept) and tested on confidence (A-D) and choice (E-H). (**I**-**P**) Cross-variable decoding using axes trained on confidence and tested on choice (I-L) and vice versa (M-P). Above chance (green) and below chance (blue) decoding indicate aligned or reversed representations, respectively (see Methods).

**Figure S7:**
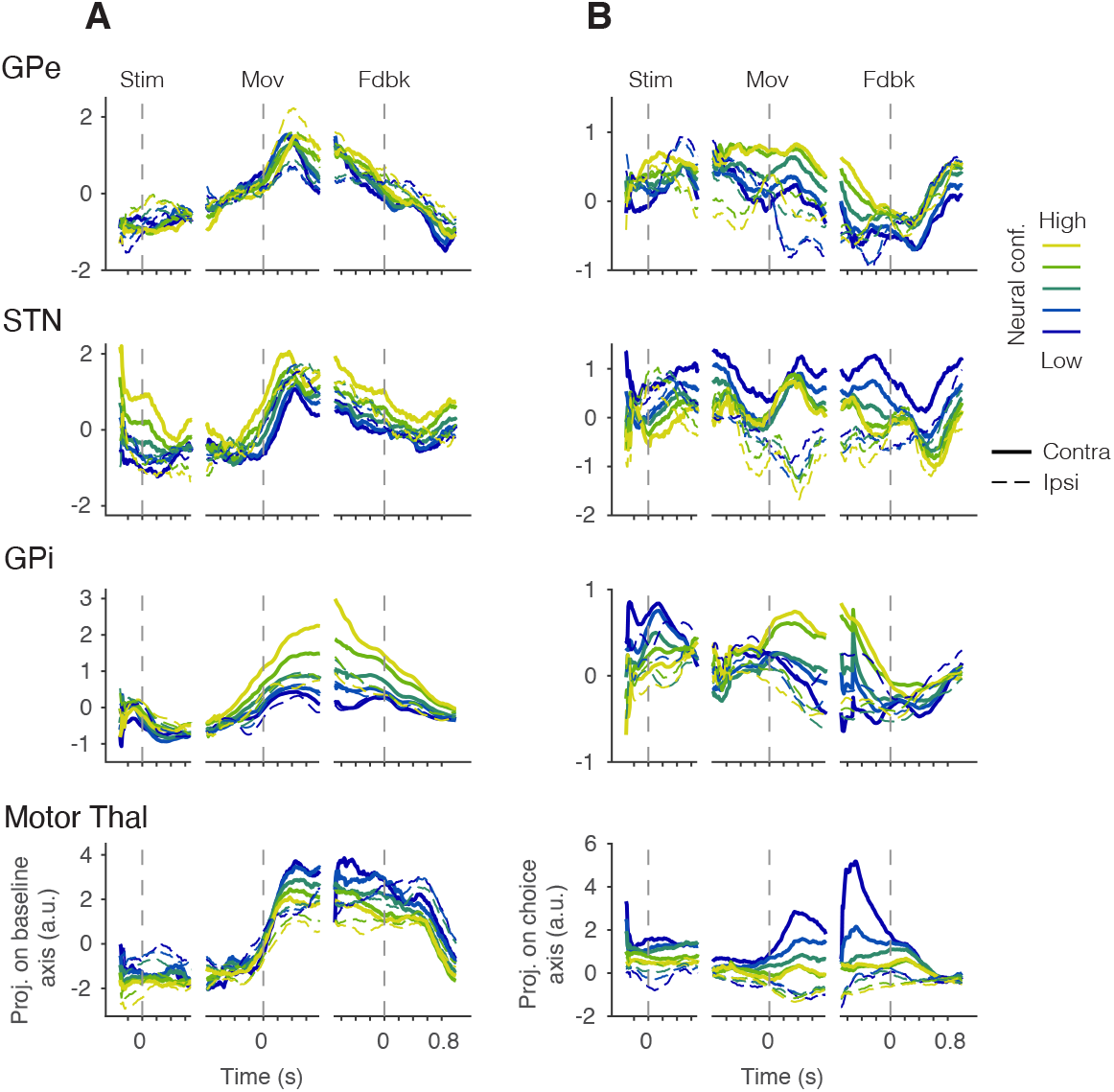
State-space trajectories of population activity. Population activity projected onto the baseline modulation axis (A) and choice axis (B). These axes are constructed from firing rates 400 ms after movement onset and are orthogonal to each other and to the confidence axis in Figure 3.

**Figure S8:**
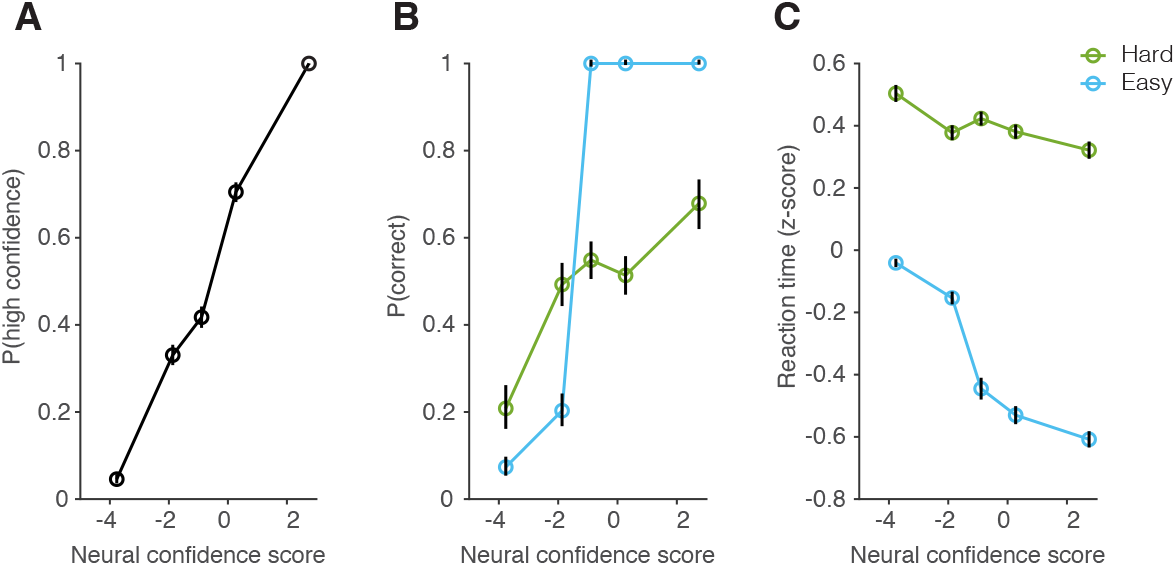
Behavioral relevance of the confidence encoding axis. Projection of motor thalamus neural activity onto the pre-feedback confidence axis predicted reported confidence (A), trial outcome (B), and reaction time (C).

**Figure S9:**
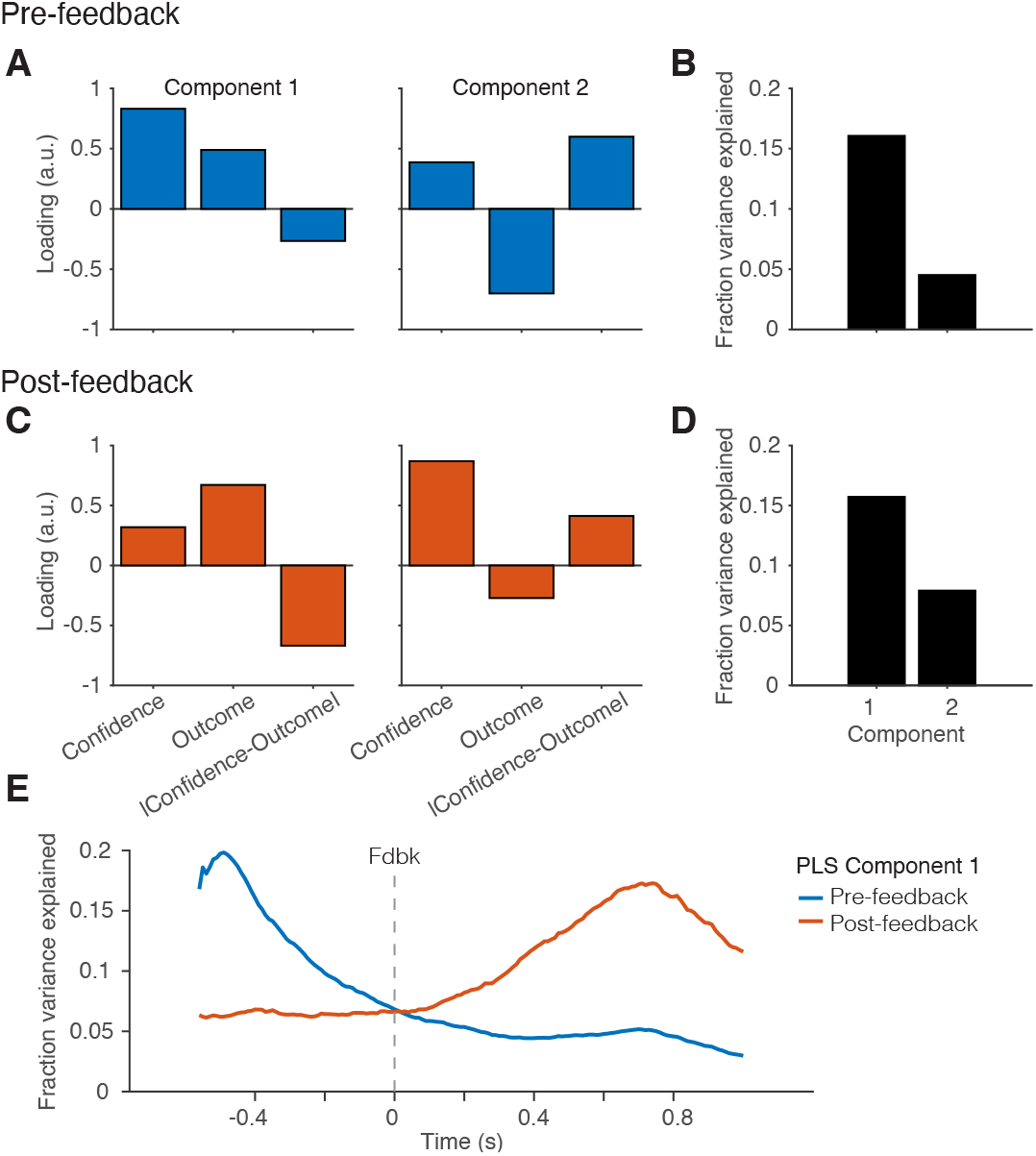
Partial least squares analysis of motor thalamus activity revealed gradual transition from encoding confidence to encoding unsigned reward prediction error. (**A**-**B**) Loadings and explained variance of task variables 400 ms before feedback. (**C**-**D**) Post-feedback PLS (600 ms after feedback). (**E**) The fraction of variance of population firing rates explained by the first component from the pre- or post-feedback PLS. Firing rates gradually transitioned from representing confidence (pre-feedback PLS component 1) to a combination of outcome and unsigned RPE (post-feedback PLS component 1).

## Notes

### Competing Interest Statement

The authors have declared no competing interest.

### Summary of Updates

We have corrected spelling errors in a few places.

